# Naturally-occurring, strain-specific defects in the retraction of *Acinetobacter baumannii* type IV pili promote biofilm formation

**DOI:** 10.1101/2024.11.14.623652

**Authors:** Yafan Yu, Rabab Mahdi, Ahmad Al-Hilfy Leon, Nam Vo, Reese Lofgren, Jean Luc Mutabazi, Kurt H. Piepenbrink

## Abstract

Type IV pili are helical filaments composed of protein subunits which are produced by numerous taxa of bacteria, including *Acinetobacter*. Type IV pili are extended out from the cell by extension enzyme complexes, which extract subunits from the membrane and insert them into the base of the filament, but can also be retracted by reverse rotation catalyzed by a retraction enzyme. Type IV pili have diverse functions, and some (twitching motility, DNA-uptake) require retraction while others (host adhesion, bacterial aggregation) do not. *Acinetobacter* bacteria, including International Clone I (IC-I) and International Clone II (IC-II) strains, show variable phenotypes in assays of type IV pilus-dependent functions. We show this variation is the result of variable efficiency in pilus retraction between pilus subtypes, and from that, a differential balance between retraction-dependent and retraction-independent functions. We define type IV pilus subtypes based on the sequence of the major subunit, PilA. In both naturally-occurring *pilA* variants from the IC-I and IC-II groups and isogenic strains complemented with IC-I or IC-II *pilA*, the IC-I pilus subtype promotes greater twitching motility and DNA-uptake while the IC-II pilus subtype promotes biofilm formation while showing reduced capacity for DNA-uptake and twitching motility, similar to a retraction-deficient mutant and consistent with the hypothesis that pilus retraction of the IC-II pilus is naturally deficient. Testing the hypothesis that this defect in retraction was sufficient to increase the level of piliation on the cell surface, we compared the yields of T4P sheared from the cell surface and found that in an isogenic background, complementation with IC-II *pilA* results in greater levels of surface PilA per cell than equivalent complementation with an IC-I *pilA* gene.

## Introduction

Type IV filaments, encompassing type IV pili (T4P) and related fibers, Flp/Tad pili and competence (com) pili are found in a wide variety of bacterial taxa and show significant variation in both structure and function (*1-5*). Type IV pili, in particular, have historically been divided into multiple subtypes; type IVa and type IVb, with many recent publications categorizing tad/flp pili as within T4P as type IVc pili (*6, 7*). While some species produce multiple T4P systems (as an example, the *Vibrio cholerae* genome contains three sets of T4P genes, producing Chitin-Regulated (ChiRP) T4Pa pili, mannose-sensitive hemagglutinin (MSHA) T4Pa pili and the T4Pb toxin-coregulated pili (*8-10*), the consequences of naturally-occurring variation within T4P systems is less well understood.

Variation in T4P subunits could result from diversifying selection with evolutionary pressure being applied by the host adaptive immune system, both antibodies and class II MHC molecules, as well as phage virions which recognize T4P (*11-15*). This variation includes the glycosylation of the major pilin subunit in both *Acinetobacter* and *Pseudomonas* (*11, 12, 16*). In addition to variation in glycosylation, natural variation in the polypeptide sequence of the major pilin of *Pseudomonas aeruginosa* appears to limit the host ranges of T4P-dependent phages (*17*). Efforts to use the major T4P subunit for vaccine development have shown substantial variation in serology leading to efforts using subunits from a range of bacterial strains (*18*) or the creation of chimeric constructs (*19*).

In the *Acinetobacter baumannii-calcoaceticus* complex (Acb), non-flagellated, Gram-negative coccobacilli associated with multidrug resistant infections in healthcare settings, type IV pili are involved in several distinct physiological processes (natural competence, twitching motility, host cell adhesion and biofilm formation) and structurally variable (*16, 20*). Previously we found structural divergence in the atomic structures of PilA, the major pilin, from three strains of *A. baumannii* (*16, 21*). We resolved the X-ray crystal structures of PilA from three *A. baumannii* strains, ACICU, AB5075 and BIDMC57, each of which shows a closer resemblance to a homologue from another species, *Dichelobacter nodosus*, than either of the three *A. baumannii* PilA proteins does to either of the other two (*16, 21-24*) (Supplementary Figure 1).

In an isogenic background, complementation of a *pilA* null mutant with these three *pilA* genes also resulted in distinct phenotypes for T4P-dependent functions, with +*pilA*^ACICU^ (from the International Clone II, IC-II group) showing very low twitching motility, but higher *in vitro* biofilm formation, than +*pilA*^AB5075^ (from the International Clone I, IC-I group) (*21*). We attributed these phenotypes to differences in surface electrostatics between the two PilA proteins, which we hypothesized would lead to greater bundling of pili between cells for the IC-II T4P. *Trans* pilus bundling (between cells) would nucleate biofilm (as observed for the bundle-forming pilus, BFP (*25, 26*)), but would reduce cellular motility by limiting the ability of cells to move independently of each other. However, observations from other bacterial species show that in some cases bundling of pili from the same cell (*cis* bundling) promotes, and may even be essential for, twitching motility (*27-29*).

One potential resolution to this conundrum is to consider DNA-uptake, another T4P function that requires retraction, the ability to depolymerase the pilus fiber, restoring the pilin subunits to the inner membrane and correspondingly shortening the pilus. The *Acinetobacter* T4P system is capable of DNA-uptake, the first step in natural competence, along with twitching motility, and the induction of biofilm formation and host-cell adherence (**Figure 1**). Conceptually, we can divide these four functions into retraction-dependent functions (twitching motility and DNA-uptake) and retraction-independent functions (biofilm formation and host-cell adherence) which are purely adhesive and independent of retraction. These differences can be observed in T4P systems through mutations of T4P-associated cytoplasmic AAA+ ATPase motors. Extension ATPases (PilB in *A. baumannii, A. baylyi* also expresses TfpB) are required to polymerize the pilins into the pilus fiber and *pilB*-null mutants, or cells where PilB is inhibited, show no T4P activity (*30, 31*). Retraction ATPases (PilT and PilU in *A. baumannii*) depolymerize the pilus fiber and *pilT*-null mutants show no twitching motility or natural competence (*20*), but the pilT mutant is hyper-piliated (*20*) and we previously found an increase in host cell adhesion in a *pilT* mutant (*16*).

**Figure 1:**
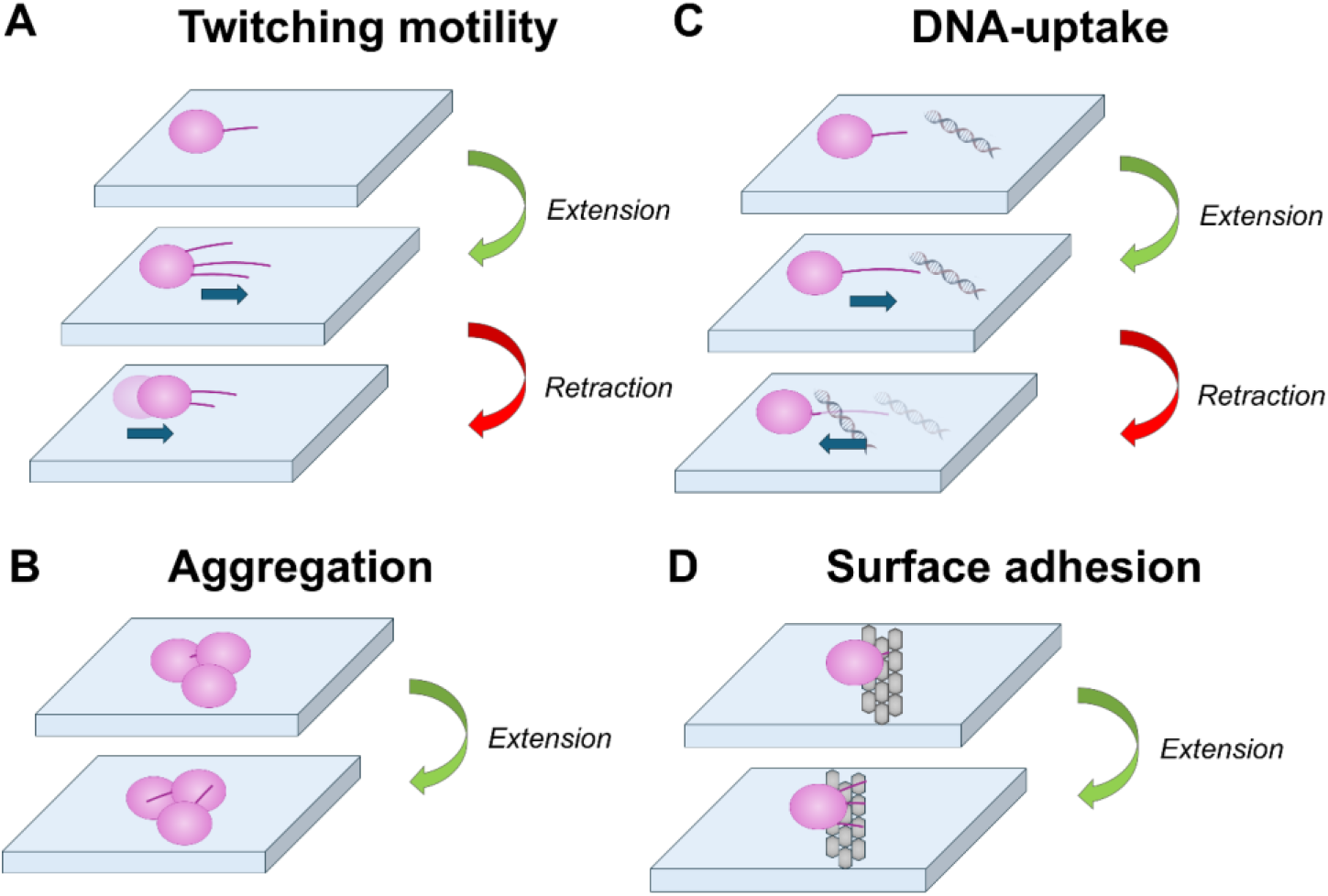
Retraction-dependent and Retraction-independent T4P activities. Schematic description of four type IV pilus functions, A) twitching motility and C) DNA-uptake (which require the ability to retract pili) and B) Cellular aggregation / biofilm formation and D) surface adhesion (which are independent of retraction).

Here, we compare retraction-depending T4P functions (DNA-uptake and twitching motility) as well as retraction-independent biofilm formation in our isogenic *pilA* complement model as well as between naturally-ocurring IC-I and IC-II *A. baumannii* strains. We found that IC-II T4P, both in an isogenic background and in naturally-occurring *A. baumanii* strains, show defects in both twitching motility and DNA-uptake, but show a greater induction of biofilm when compared to their IC-I counterparts which may be explained by greater surface T4P expression. Both of these features, loss of retraction-dependent T4P functions and gain of biofilm formation, appear to be the consequences of a natural defect in T4P-retraction resulting from the IC-II T4P structure.

## Materials and Methods

### Strains and plasmids

Details on *A. nosocomialis* M2 and mutants can be found here (*20*) (where it was originally identified as *A. baumannii* M2). *A. baumannii* AB5075 wild type and mutants are from the University of Washington AB5075-UW transposon mutant library maintained by the Manoil group (*32*). Strains from the International Clone I and International Clone II groups were provided by the American Type Culture Collection (ATCC). Details of complementation *by pilA* genes from *A. baumannii* AB5075, ACICU and BIDMC57 can be found in Ronish *et al*. (*21*). The *pilA* ACICU negative and αβ-loop mutants (sequences available in Supplementary Table 1) were synthesized by Genscript and ligated into pUCP20GM (*33*) using BamH1 and HindIII restriction sites. The resulting vectors were electroporated into *A. nosocomialis* M2 Δ*pilA* (*20*) using standard protocols (*34*). The presence of the plasmids was confirmed by both resistance to gentamycin and PCR of the pilin genes. For plasmid maintenance, strains with strains complemented with *pilA*::pUCP20GM were grown on solid media supplemented with 50ug/ml gentamycin before their use in the assays described below.

### Twitching motility

Macroscopic twitching motility was assessed using standard protocols (*16, 21, 35*). Briefly, *Acinetobacter* strains, including mutants and complemented strains, were grown on 1.5% MacConkey agar plates overnight. Colonies were selected and stabbed through the centers of 1% MacConkey agar plates in polystyrene petri dishes. The plates were incubated in sealed bags at 37°C for three days. The agar was then removed, and the bacteria adhered to the polystyrene petri dish stained with 0.1% crystal violet for 5 minutes. Excess crystal violet was removed by gentle washing with DI water 3 times. The subsurface twitching area on each plate was accessed by measuring the diameters of the ellipse and then calculating the area (*36*). Statistics were calculated for three replicates and significance determined by Student’s t-test.

### Static biofilm formation

Static biofilm formation was quantified using horizontal growth on 6-well plates, with frequent changes of media and quantification through staining with crystal violet. Briefly, *Acinetobacter* strains, including mutants and complemented strains, were grown in LB no salt broth (Tryptone 10 g/L, Yeast Extract 5 g/L) overnight. The bacteria were inoculated into 6-well plates to have a final OD600 of 0.1 in 5ml total volume. The 6-well plates were incubated at 37°C for 3 days (72h). To remove non-adherent, planktonic bacteria and introduce fresh growth media, the medium was changed every 20-24h using pipette. After 72h, 5ml of PBS was added to and removed from the plates to wash the planktonic bacteria. 5ml of 0.1% crystal violet with 2.5% glutaraldehyde was added to the plates and stained for 10min. Excess crystal violet was removed by gentle washing with PBS. 1 ml of 100% ethanol was then added to the plates to dissolve the dye. Relative biomass was assessed using the final concentration of released crystal violet; the absorbance value was read using a NanoDrop-C at 570 nm.

### DNA-uptake through flow cytometry

Uptake of extracellular DNA was quantified using flow cytometry to measure the proportion of fluorescent bacterial cells after incubation with a plasmid encoding mCherry. This method is adapted from Godeux *et al*. (*37*). Briefly, *Acinetobacter* strains, including mutants and complemented strains, were grown in LB no salt broth overnight. The cultures were then diluted 1:200 into 200ul LB no salt broth, and 500ng of plasmid encoding mCherry was added. The broth was then grown at 37°C for 4h. Bacterial suspensions were fixed with formaldehyde (final concentration, 3.7%) for 30 min at room temperature and then washed twice with PBS and further resuspended in 200ul PBS. Flow cytometry was performed using a Beckman Coulter CytoFLEX LX. Samples were run at a collection rate of 60ul/min, the mCherry was excited by the 561nm laser and fluorescence emission was detected using a 610/20 band pass filter. Data was acquired using CytExpert softare (Beckman Coulter), and further analyzer in FlowJo (BD Biosciences).

### Quantifying pilus expression

Pilin proteins were isolated using the methods of Voisin *et al*. (*38*) with modifications. Bacteria were spread on ten LB no salt agar plates and grown at 30°C overnight. Next day the bacteria were gently scraped from each plate using a sterile cell scraper and resuspended in a 3 ml buffer containing 50 mM Tris-HCl and 150 mM NaCl (pH 8.3). The cells were injected through needle (diameter 18G 1” Blunt needle+ 25G 1”) three times to shear off the pili. The cell suspension was then spin down for 5 minutes at 13300 x g at 4°C. The supernatant was transferred to new 1.5 ml microcentrifuge tubes and centrifuged for an additional 25 minutes for same speed at 4°C. For precipitating the sheared pili, 1M MgCl2 was added to the supernatant to a final concentration of 0.1M and the samples were incubated at 4°C overnight. Samples were further centrifuged for 25 minutes at 13300 x g at 4°C. Then the pellet was resuspended in 100 µl buffer containing 50 mM Tris-HCl and 150 mM NaCl (pH 8.3) and stored at 4°C. Relative PilA surface expression was compared using densitometry by ImageJ from silver-stained SDS-page gels (*36*).

### Figures and diagrams

Data figures were created using GraphPad Prism. Images of the PilA^ACICU^ (PDB ID: 4XA2) and PilA^AB5075^ (PDB ID: 5VAW) X-ray crystal structures were made using UCSF ChimeraX (*39*) and the dendrogram of PilA sequences was created using Interactive Tree of Life (*40*).

### Statistical analysis

All statistical analyses were performed using GraphPad Prism version 9.5.0 (GraphPad Software, San Diego, California USA). All significant differences were determined at p-value<0.05. Relevant statistical tests used for comparisons are discussed in figure captions.

## Results

### In an isogenic background, IC-II T4P show decreased competence compared to IC-I T4P

Natural competence or natural transformation, the ability to take up, internalize and replicate extracellular DNA, is directly reliant on DNA-uptake and previously, mutants of the T4P system abrogating either pilus biogenesis (*41*), pilus retraction (*20*), or DNA binding (*42*) have been shown to have defects in natural competence. To compare the relative ability of IC-I and IC-II T4P systems to take up extracellular DNA, we used isogenic chimeric strains expressing *pilA* from strains AB5075 (IC-I) or ACICU (IC-II) using the pUCP20GM plasmid as previously described (*21*). All strains were grown to exponential phase and incubated with an mCherry plasmid in broth. Uptake of the plasmid (and hence red fluorescence from the production of mCherry) was quantified for each strain using flow cytometry as described above, with methods adapted from Godeux *et al*. (*37*). Results from this experiment are shown in Figure 2 with panel A showing representative data from the +*pilA*^AB5075^ and +*pilA*^ACICU^ complemented strains while panel B shows the complete cross-strain comparison. As expected, wild type *Acinetobacter nosocomialis* M2 bacteria (the parent strain) are capable of natural competence under these conditions, but Δ*pilA* and Δ*pilT* strains which are deficient in retraction (Δ*pilT*) or pilus biogenesis (Δ*pilA*), showed no natural competence. For the complemented strains, we found that the AB5075 complement was competent, but the ACICU complement was significantly less able to take up DNA (p = 0.0078), statistically indistinguishable from the Δ*pilA* strain it was derived from (p = 0.3739).

**Figure 2:**
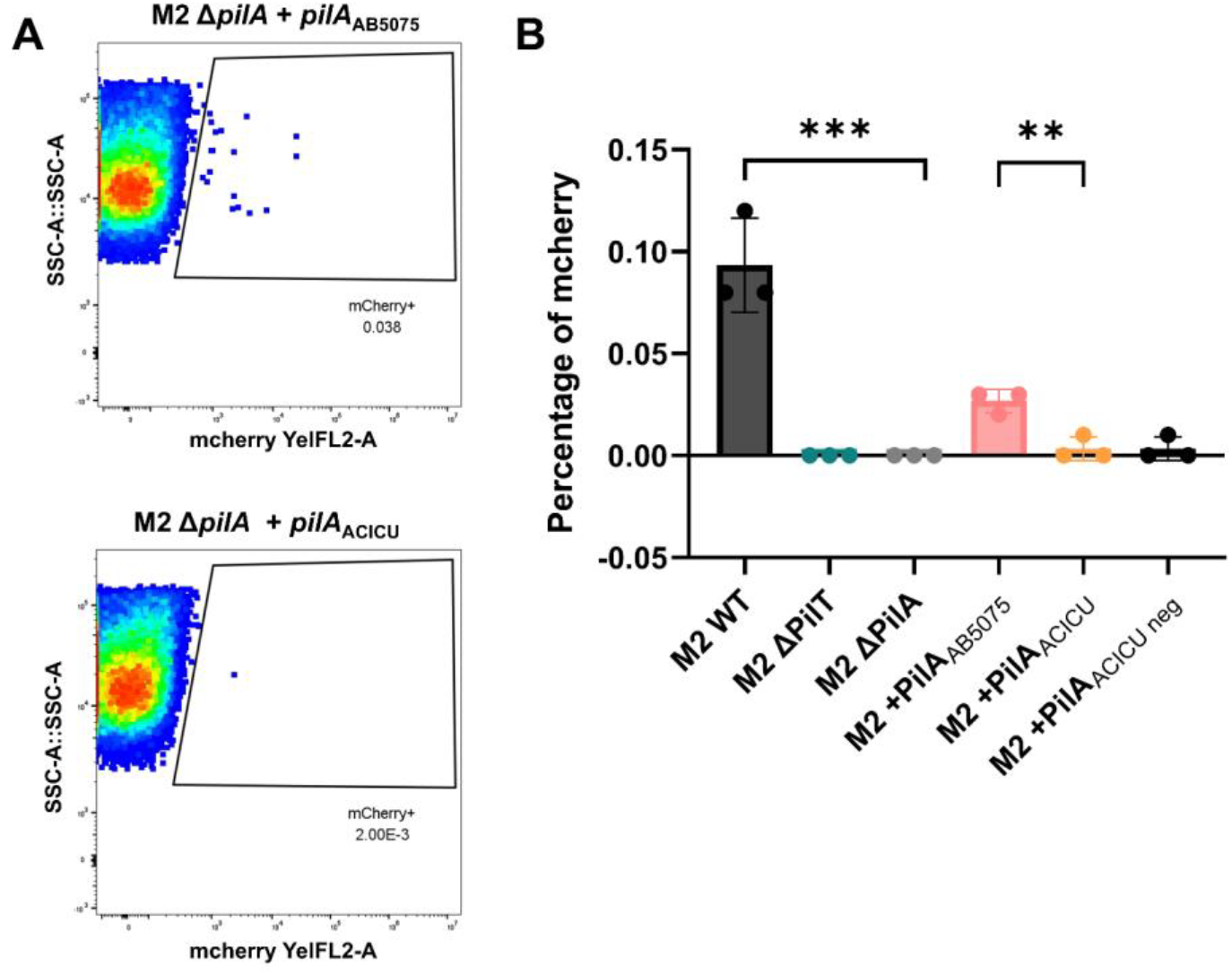
DNA-uptake by IC-I and IC-II T4P in an Isogenic Background. A) representative data from DNA-uptake measurements (of an mCherry-encoding plasmid) by flow cytometry for *A. nosocomialis* M2 Δ*pilA* + *pilA*^AB5075^ (IC-I) and *A. nosocomialis* M2 Δ*pilA* + *pilA*^ACICU^ (IC-II). B) DNA-uptake for M2 wt, mutants and complements. Significant differences are measured by ANOVA, * p ≤ 0.05, ** p ≤ 0.01, *** p ≤ 0001, **** p ≤ 0.0001

### Differences in surface electrostatics cannot explain differential T4P phenotypes

Previously, we found that in this isogenic background, bacteria with AB5075 (IC-I) T4P showed higher twitching motility and lower biofilm formation than those with ACICU (IC-II) T4P (*21*). We also found that, based on X-ray crystal structures of PilA^ACICU^ and PilA^AB5075^, the AB5075 pilus would have a more electronegative surface area than its ACICU counterpart. We hypothesized a structure-function relationship where T4P-bundling between adjacent cells would promote biofilm formation (by nucleating assembly) and hinder motility (by preventing these cells from moving independently) and that the relatively electronegative AB5075 T4P fibers would be less prone to bundling. To test this hypothesis, we made two chimeric mutant *pilA* genes based on structural differences between PilA^ACICU^ and PilA^AB5075^, highlighted in Figure 3, the αβ-loop (panel A) and five surface-exposed basic residues found in PilA^ACICU^ (panel B). Because the electronegativity of the PilA^AB5075^ surface was driven not by an over-abundance of acidic residues, but the absence of basic residues on the surface, we created a *pilA*^ACICU^ gene where each of the surface exposed, basic residues was mutated to the counterpart residues in PilA^AB5075^, K102T, K103T, R122T, K124E, R132N, *pilA*^ACICU-negative^. Because the greatest concentration of structural differences between the two PilA proteins is in the αβ-loop, we also created a mutant *pilA*^ACICU^ gene where the sequence of the PilA^AB5075^ αβ-loop (residues 54-93) was substituted for the PilA^ACICU^ αβ-loop (residues 54-78), *pilA*^ACICU-loop^.

**Figure 3:**
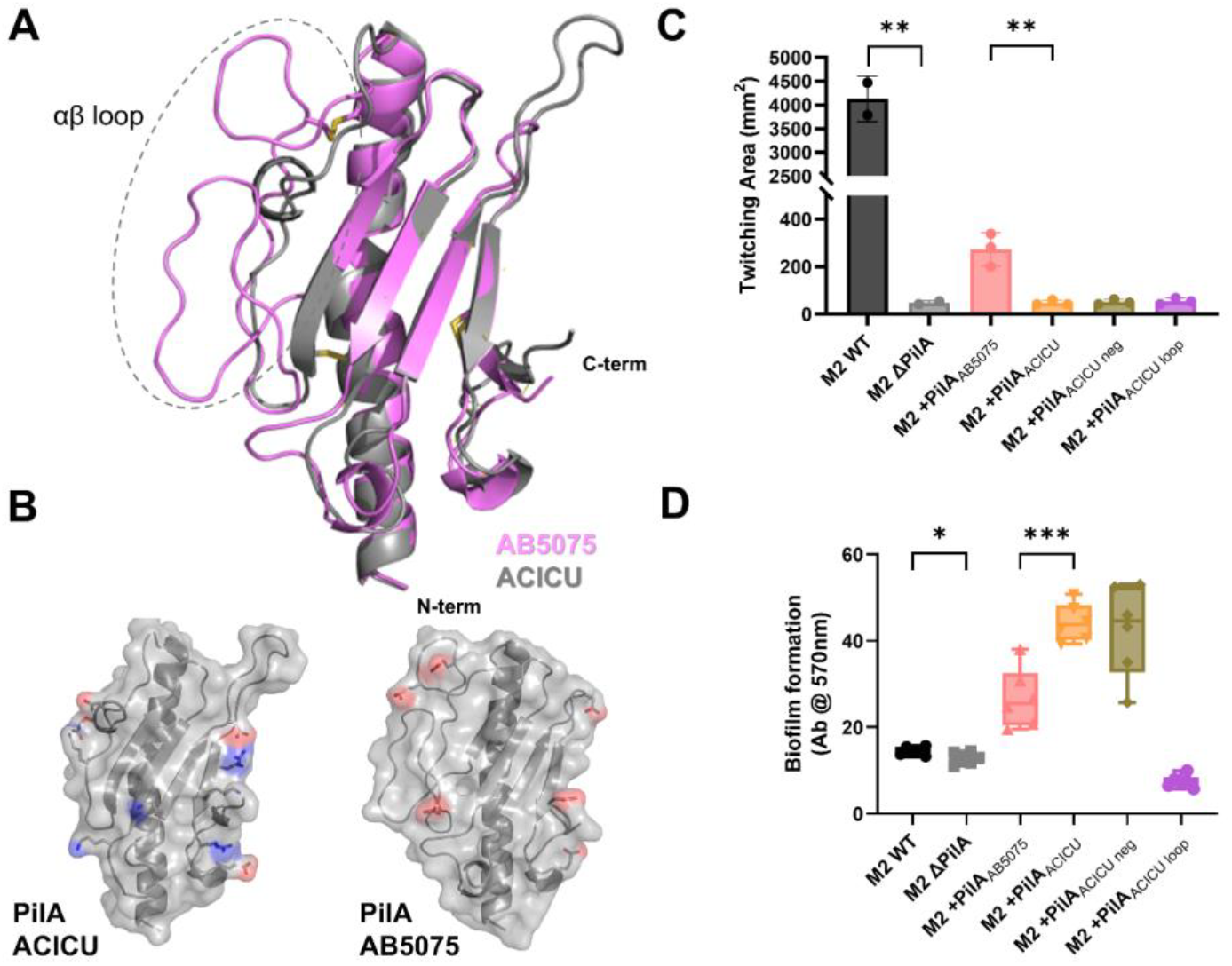
Twitching motility and Biofilm formation for PilA ACICU mutants. A) Overlay of the PilA^ACICU^ and PilA^AB5075^ structures, AB5075 is in pink with ACICU in grey with the αβ-loop encased in a dashed semicircle. B) surface charged residues PilA^ACICU^ and PilA^AB5075^ (exposed in a model of the assembled pilus) are depicted either in red (acidic, electronegative) or blue (basic, electropositive) C) twitching motility for M2 wt, Δ*pilA* and complements, D) static biofilm formation for M2 wt, Δ*pilA* and complements. Significant differences are measured by ANOVA, * p ≤ 0.05, ** p ≤ 0.01, *** p ≤ 0001, **** p ≤ 0.0001

However, as the results of functional comparisons show in Figure 3, panels C and D, neither of these alterations resulted in the high-motility, low-biofilm phenotype seen from PilA^AB5075^. Complementation with *pilA*^ACICU-loop^ resulted in a *pilA*-null phenotype for both twitching motility and biofilm formation (Figure 3) suggesting that the chimeric protein was unable to fold efficiently. The *pilA*^ACICU-negative^ complement produced results identical to *pilA*^ACICU^ in DNA-uptake (Figure 2), twitching motility (Figure 3, panel C) and biofilm formation (Figure 3, panel D). These results suggest that the difference between the AB5075 and ACICU T4P phenotypes is not a product of differential surface electrostatics, but some other differential aspect of the two pilus structures.

### International Clone II strains show defects in retraction-dependent T4P activity

Based on our findings for +*pilA*^ACICU^ and +*pilA*^AB5075^ complements in the M2 background, we hypothesized that the structural differences in IC-I and IC-II PilA proteins were responsible for the differential phenotypes for T4P functions. However, while the *pilA* gene is more variable in *Acinetobacter* than other T4P genes (*16, 21*), differences in either other genes, including other pilin subunits or pilus biogenesis genes could also impact T4P phenotypes. Prior studies by other groups have shown differences between IC-I and IC-II *A. baumannii* strains in surface behavior, particularly twitching motility (*43, 44*). To quantitatively compare T4P functions in naturally occurring strains, we obtained eight *A. baumannii* strains from the ATCC, including two IC-I and four IC-II strains. These strains (along with *A. baumanii* AB5075) were assayed for three T4P-dependent functions, twitching motility (panel B), DNA-uptake (panel C) and biofilm formation (panel D) as shown in Figure 4.

**Figure 4:**
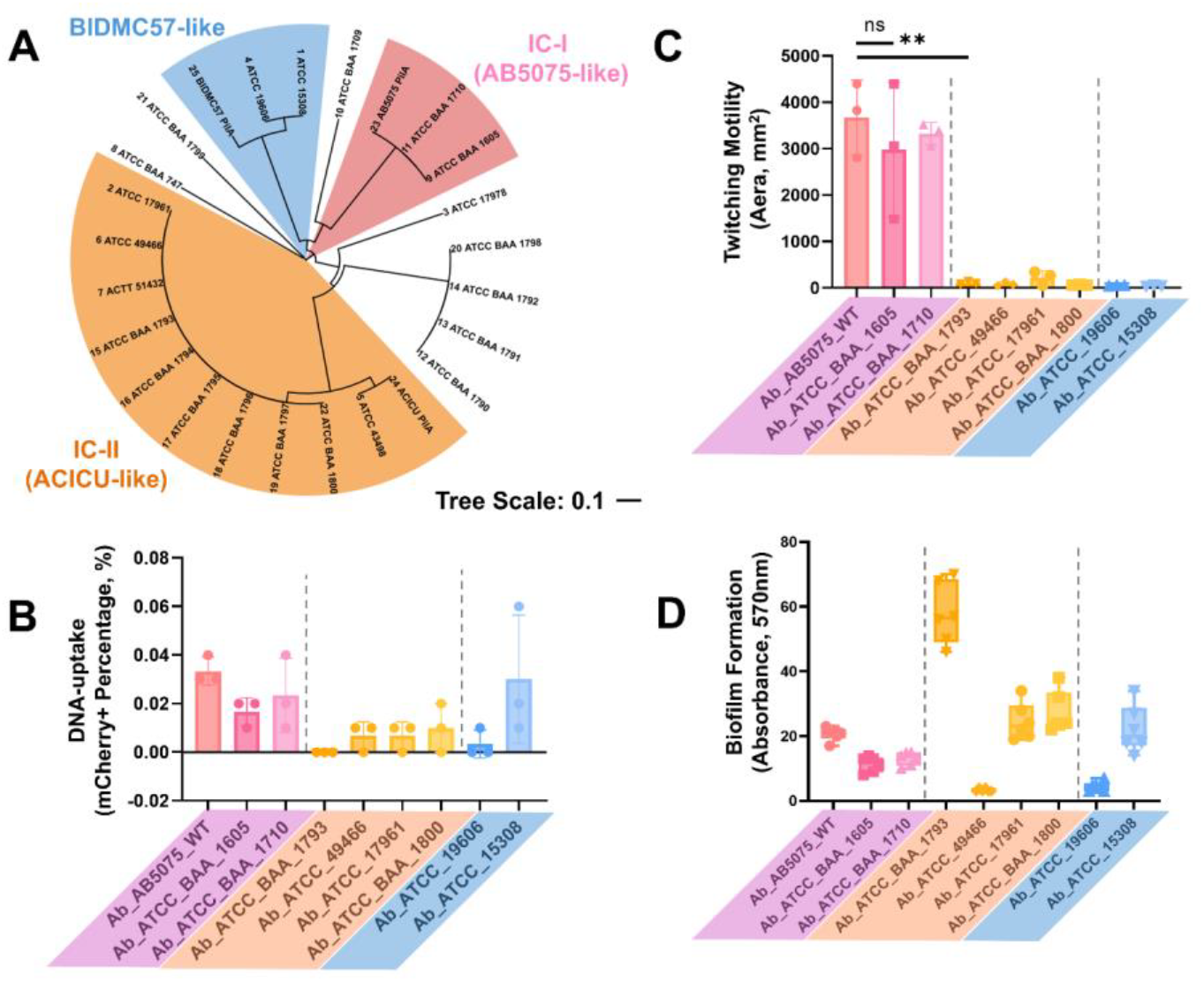
T4P-dependent Function in IC-I and IC-II *A. baumannii* strains. A) Dendrogram of *A. baumannii* strains by PilA sequence, clusters are highlighted corresponding to PilA^ACICU^ (orange), PilA^AB5075^ (pink) and PilA^BIDMC57^ (blue), B) DNA-uptake of an mCherry plasmid by flow cytometry; strains are colored by PilA subtype with the IC-I (AB5075) cluster in pinks, the IC-II (ACICU) cluster in yellows and the BIDMC-57 cluster in blues. C) twitching motility with identical coloration to panel B, and D) static biofilm formation measured by crystal violet staining with strains colored identically to panels B and C. Significant differences are measured by ANOVA, * p ≤ 0.05, ** p ≤ 0.01, *** p ≤ 0001, **** p ≤ 0.0001

All four IC-II isolates were defective for retraction, as evidenced by lack of both DNA-uptake and motility. In contrast, the IC-I strains all showed robust twitching motility and DNA-uptake. The two isolates from the third group (with BIDMC-57-like PilA proteins) were non-motile but ATCC 15308 showed robust DNA-uptake and modest biofilm formation, similar to the IC-I strains. On average, IC-II isolates formed more biofilm than IC-I isolates, but ATCC 49466 showed a defect for biofilm formation, similar to *pilA* mutant. We hypothesize that ATCC 49466, along with ATCC 19606 are largely unable to generate T4P under these experimental conditions as they were negative for all three functions. This observation is consistent with negative results for ATCC 19606 twitching motility observed by others (*43, 45*) despite the fact that PilA in *A. baumannii* ATCC 19606 and PilA in *A. nosocomialis* M2 are 92% identical (Supplementary Figure 2) and M2 shows twitching motility under identical conditions (Figure 3, panel C).

To evaluate the impact of retraction deficiency on biofilm formation, we measured static biofilm formation for this panel of nine naturally occurring *Acinetobacter baumannii* strains. As expected, IC-II (ACICU-like) strains showed enhanced biofilm formation, on the average, compared to IC-I (AB5075-like) strains (Figure 4, panel D). Notably, despite belonging to IC-II, *A. baumannii* ATCC 49466 exhibited reduced biofilm formation (along with poor twitching motility and DNA-uptake), likely due to incompatibility with the growth conditions used in these assays, similar to the results described above for *A. baumannii* ATCC 19606. These findings suggest that biofilm formation does not require T4P retraction, and in fact, retraction deficiency promotes biofilm formation in *Acinetobacter baumannii*.

### Retraction-deficiency induces biofilm formation in Acinetobacter baumannii

To test the hypothesis that retraction deficiency promotes retraction-independent T4P functions, including biofilm formation, we measured static biofilm formation in *A. nosocomialis* M2 and *A. baumannii* AB5075 for the wild type strains and *pilT* mutants (M2 Δ*pilT* and AB5075 Tn-*pilT*-). Prior studies in multiple organisms have shown that *pilT* is required for efficient retraction of T4P as evidenced by DNA-uptake/natural transformation and macroscopic twitching motility (*20, 46, 47*). Our group previously found that an M2 *pilT* mutant showed a defect in twitching motility, but showed significantly greater adherence to A549 and Detroit 562 monolayers (*21*). Figure 5 shows the results of assays of biofilm formation (panel B) and DNA-uptake (panel C). Regardless of genetic background, the *pilT*-mutants showed defects in DNA-uptake, but an induction of static biofilm compared to the parent strains. Unlike comparisons between IC-I and IC-II *A. baumannii* strains or the *pilA*^AB5075^ and *pilA*^ACICU^ complements of M2 Δ*pilA*, these effects can be ascribed solely to defects in retraction as no modification was made to the *pilA* genes. We hypothesize that this induction in biofilm formation is driven by an increase in surface piliation (*33*), consistent with results from isolation of sheared pili from the surface of *A. baumannii* cells as described below.

**Figure 5:**
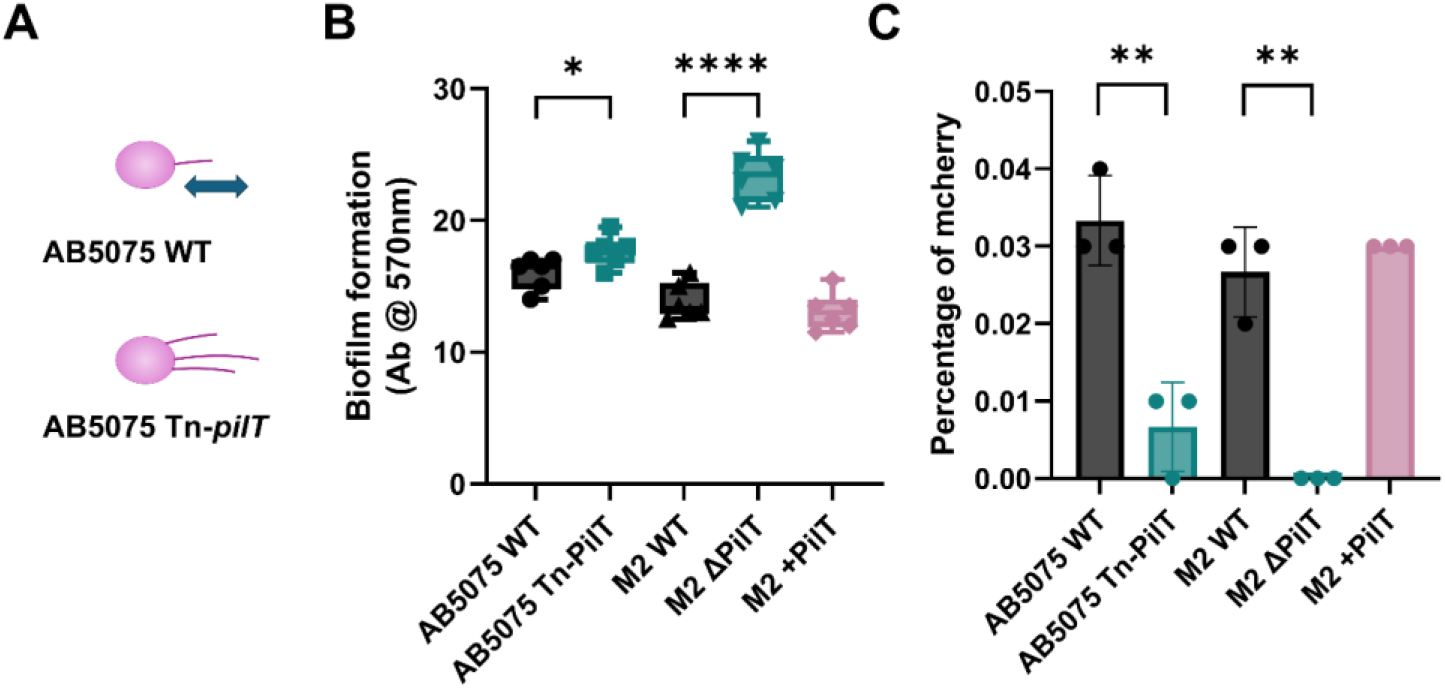
DNA-uptake and Biofilm Formation by *pilT* Mutants. A) schematic representation of *A. baumannii* wild type and Δ*pilT* showing the retraction defect and increase in piliation for Δ*pilT*, B) static biofilm formation for *A. nosocomialis* M2 and *A. baumannii* AB5075 *pilT* mutants, C) DNA-uptake by flow cytometry for *pilT* mutants. Significant differences are measured by ANOVA, * p ≤ 0.05, ** p ≤ 0.01, *** p ≤ 0001, **** p ≤ 0.0001

### Complementation with IC-II T4P leads to greater surface expression of T4P

If defects in retraction can explain the greater induction of *in vitro* biofilm formation shown by *pilT* mutants and IC-II T4P systems, we would expect both to be products of greater surface expression of T4P, either in the number of pili per cell or through longer pili. In a comparison of Transmission Electron Microscopy (TEM) images, greater numbers of pili were observed previously in *A. nosocomialis* for a *pilT* mutant (*20*), but unambiguously identifying T4P (as opposed to chaperone-usher (CU) pili) is difficult at these resolutions because of their similar width, 6-8 nm (*12*).

To compare T4P surface expression between the IC-I and IC-II subtypes, we used growth conditions expected to favor the production of T4P over CU pili, selective precipitation to reduce levels of contaminants and compared levels of surface PilA by gel electrophoresis and densitometry. Comparisons of *A. baumannii* AB5075 wild type and Tn-*pilT*-showed greater surface PilA expression for the *pilT* mutant (Supplementary Figure 3). We compared *A. nosocomialis* M2 Δ*pilA* +*pilA*^ACICU^ and M2 Δ*pilA* +*pilA*^AB5075^, using the parent strain, M2 Δ*pilA* as a negative control to ensure that we could unambiguously assign the PilA band (Supplementary Figure 4). The results of this experiment are shown in Figure 6 panel A, where complementation with the IC-II subtype (*pilA*^ACICU^) resulted in significantly higher levels of PilA sheared from the surface per bacterial cell.

**Figure 6:**
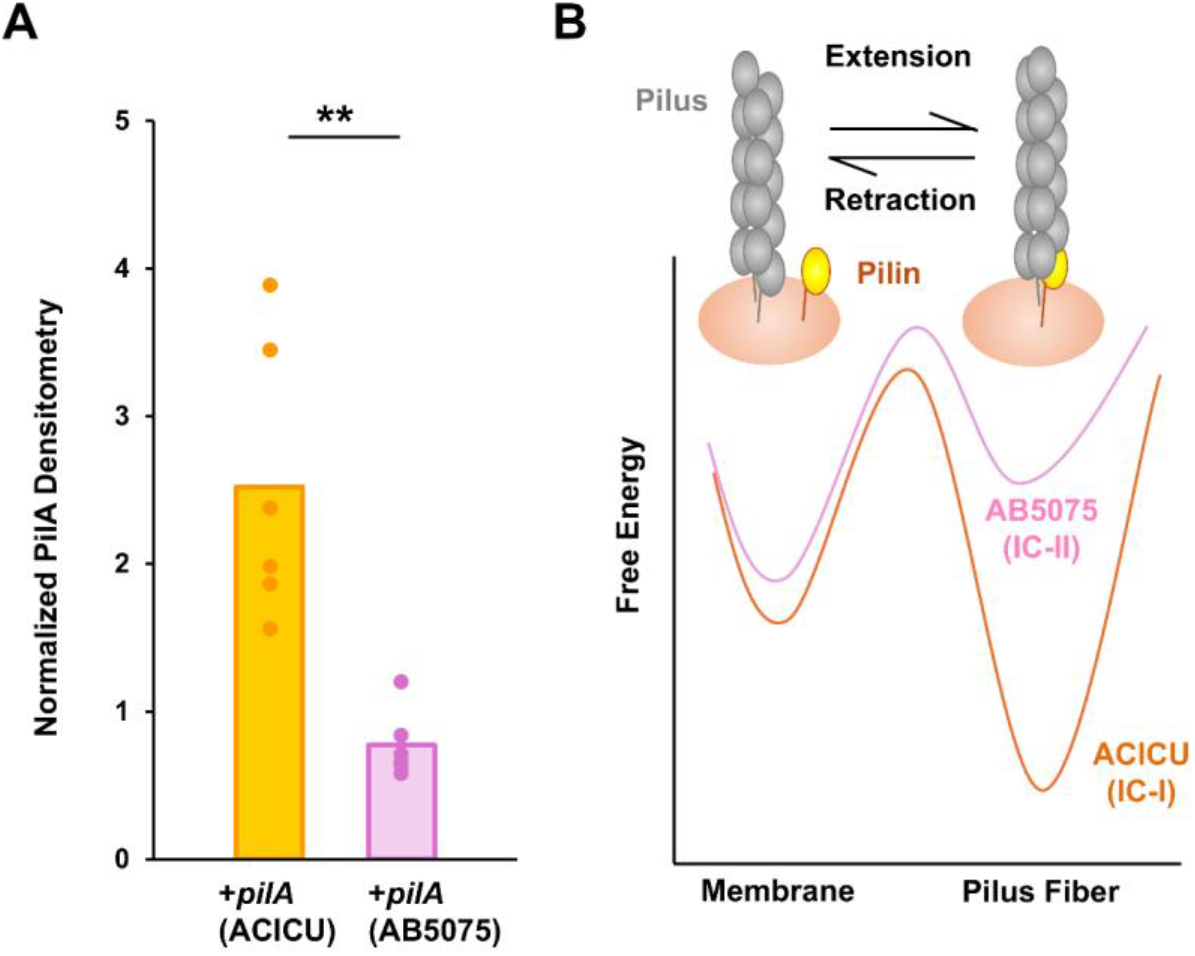
T4P Surface Expression. A) Normalized PilA surface expression for *A. nosocomialis* M2 Δ*pilA* + *pilA*^AB5075^ (IC-I) and *A. nosocomialis* M2 Δ*pilA* + *pilA*^ACICU^ (IC-II) as measured by densitometry of a PilA band using silver-staining, D) Free energy diagram showing energetic barriers for T4P retraction and the putative differences between AB5075 and ACICU T4P. Significant differences are measured from the wild type strain using Student’s one-tailed T-test, * p ≤ 0.05, ** p ≤ 0.01, *** p ≤ 0001, **** p ≤ 0.0001

## Discussion

The type IV filament structure is unique in its reversible assembly. Unlike chaperone-usher or sortase pilus systems, type IV pilins can be polymerized into pili and pili depolymerized into pilins through reversed rotations of cytoplasmic ATPase motors (PilB, PilT, PilU) attached to the same biogenesis machinery (PilC, PilM, PilN, PilO, PilP). Pilus retraction provides unique benefits for T4P systems by enabling twitching motility and DNA-uptake, which in *Acinetobacter*, are absolutely dependent upon T4P function. This stands in contrast to retraction-independent T4P adhesive activity promoting host cell adhesion and biofilm formation which occur through multiple molecular mechanisms.

Despite the obvious benefits of efficient pilus retraction, data from retraction ATPase mutants indicates that it imposes costs as well. Mutants of *pilT* in *Pseudomonas aeruginosa* were observed previously to be hyperpiliated (*33*), and Harding *et al*. were able to observe T4P on the surface of an *A. nosocomialis* M2 Δ*pilT* mutant, but not the parent strain (*20*). To the extent to which T4P act as adhesins, reductions in their number will result in decreased adhesion (and vice versa) and indeed, we show here that *Acinetobacter pilT* mutants show an induction of biofilm compared with their parent strains.

If pilus retraction provides novel functions for T4P but also restricts adhesive functions, this implies that i) T4P retraction should be viewed as a scalable quality and that an optimum likely exists between the most extreme possible cases (e.g. zero retraction or retraction outstripping extension to completely remove T4P from the cell surface) and ii) that the optimum level of retraction is likely to vary depending on the circumstances or conditions under which T4P are being produced.

As we have noted previously, the *pilA* gene in *Acinetobacter* is more variable, not only than the pilus processing/extension/retraction machinery (PilBCD, PilMNOP, etc) but other pilin genes (*16, 21*). Variation in surface-exposed residues may be driven by diversifying selection (*48*), but the variation in *Acinetobacter* PilA is not limited to surface exposed residues but includes substantial differences in regions likely to impact pilus assembly. Taken together, these features imply that variation in *pilA* may be a critical mechanism for modulating T4P functions, not only in *Acinetobacter baumannii*, but in related species which are found both in the environment and as opportunistic pathogens and where the major T4P subunit shows similar variation, including *P. aeruginosa* and *D. nodosus*. However, we note that *P. aeruginosa* strains with PilA subunits structurally similar to PilA^ACICU^, PAO1 and PAK are more motile than *A. baumannii* IC-II strains, suggesting that other factors may also impact the balance between extension and retraction.

Here, we show that variation in surface-exposed residues between two T4P subtypes, PilA^ACICU^ and PilA^AB5075^, has no direct impact on the functional differences we observe both in an isogenic background and in the naturally-occurring IC-I and IC-II strains, as evidenced by the commonality of phenotypes between the M2 Δ*pilA* +*pilA*^ACICU^, *pilA*^ACICU_negative^ and naturally-occurring IC-II bacteria. Instead, differences between the two are driven by a combination of decreased pilus retraction and increased surface piliation for the IC-II T4P system, both of which could be explained by increased pilus stability creating an energetic barrier to pilus retraction.

Clinical epidemiology also suggests that *Acinetobacter* isolates vary in T4P-mediated surface behavior based on their origin, with lung isolates showing, in aggregate, greater biofilm formation and blood isolates greater twitching motility (*49*). We hypothesize that these differences stem from the distribution of T4P subtypes within these populations. In a survey of *Acinetobacter* infections in Japan, Matsui *et al*. found that IC-II strains accounted for 43% of *A. baumannii* respiratory infections but none of the blood infections and only 2% of all other infections (*50*). Comparing surface behavior from different *A. baumannii* isolates, Eijkelkamp *et al*. characterized only 9% of IC-II isolates as positive for twitching motility, but 100% of IC-I isolates were positive. Conversely, median biofilm formation (as measured by crystal violet) was found to be approximately 70% greater for IC-II isolates (*43*). Skerniškytė *et al*. also found greater twitching (but not swarming) motility for IC-I strains compared to IC-II and greater desiccation-resistance for IC-II strains (*44*). Vijayakumar *et al*. also found that that *A. baumannii* isolates varied considerably in their surface behavior based on the site of isolation. Their results showed that respiratory isolates formed more biofilm but were less motile in twitching assays than blood isolates (*49*). We believe that all of these results can be explained through differences in pilus retraction similar to those observed here.

Despite the strong association in the literature between T4P and twitching motility (*20, 27, 29, 42, 46, 51-57*), the evolution of retraction-deficient T4P systems in *Acinetobacter* is plausible not only because of the potential advantages for adhesive functions, but because pilus retraction may be more useful in environmental than in host lifecycles and may carry associated risks. Although mutants of *pilA* have been found to be less pathogenic in an animal model (*58*), twitching motility itself appears to be less important than surface motility (*59*) *in vivo* and IC-II strains with poor motility are still virulent in murine infections (*44*). Pilus retraction also carries risks in terms of phage infection as retraction is an essential step in T4P-mediated phage infection and *pilT* mutants are immune (*11, 13*).

If *A. baumannii* T4P systems are, in fact, specialized for distinct functions, that specialization must stem from measurable physical differences between some component(s) of pilus itself or the apparatus responsible for pilus extension and retraction. Here we show that we can reproduce the phenotypic differences found in naturally-occurring IC-I and IC-II strains in an isogenic background solely through modifications in *pilA*, In *A. baumannii*, there are over a dozen gene-products necessary for pilus biogenesis and function, including seven putative pilin subunits, but it is nevertheless unsurprising that PilA, which makes up the vast majority of the pilus, has a controlling role. Because the vast majority of the subunits in a type IV pilus are PilA, interactions between the pilus machinery during extension and retraction are primarily with PilA, and interactions between PilA subunits are the principal stabilizing force of pilus assembly. In Figure 6, panel B, we present a model by which an increase in stabilizing interactions between assembled PilA subunits could create an energetic barrier to retraction which decreases retraction speed by opposing the force of the PilT motor. We believe this model represents the best current explanation for the phenotypic differences in T4P function, both because, as we have shown here, these differences can reproduced solely through variation in *pilA* in an isogenic background and because complementation with an IC-II *pilA* gene produces relative defects in both retraction-mediated T4P functions when compared to an IC-I *pilA* complement.

Taken together, the results here suggest that the evolution of pilus retraction is a double-edged sword, with the addition of new capabilities (twitching motility and DNA-uptake) balanced against a reduction in adhesive capability. This balancing act may favor different pilus structures with different levels of potential retraction depending on the conditions in the host or other environment and suggests that variation on the structures of T4P subunits within bacterial species is partially a product of lifecycle adaptation.

## Acknowledgements

We wish to acknowledge outstanding technical assistance by Dirk Anderson at the Nebraska Center for Biotechnology Flow Cytometry Core in our DNA-uptake assays. and the Manoil group at the University of Washington for strain from the *A. baumannii* AB5075-UW mutant library.

