## Supplemental Material for "Naturally-occurring, strain-specific defects in the retraction of *Acinetobacter baumannii* type IV pili promote biofilm formation"

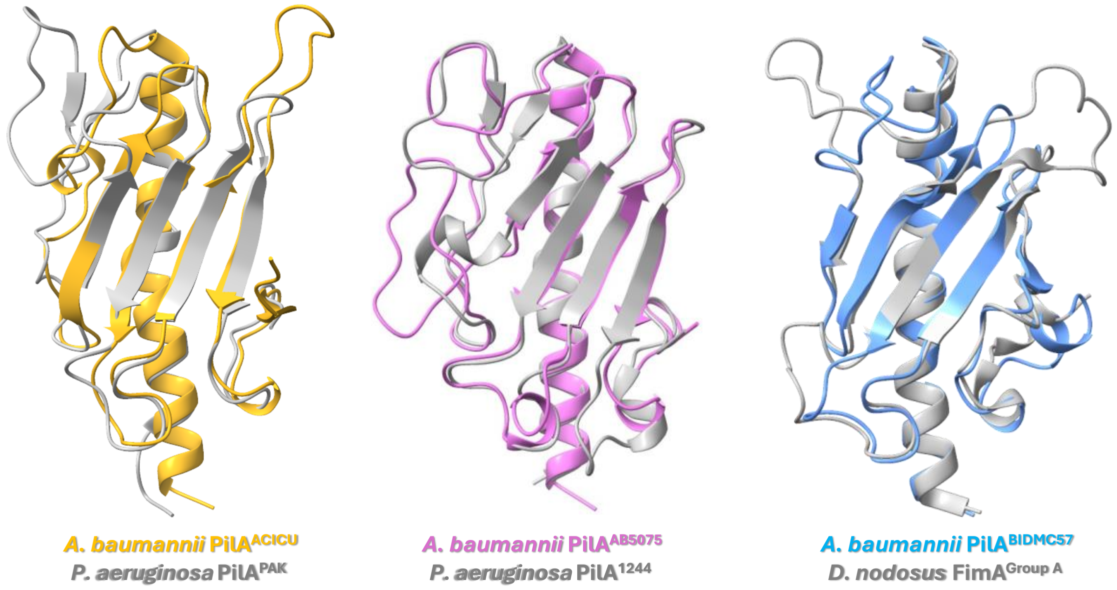


**Supplementary Figure 1:** Superimpositions of the X-ray crystal structures of pilin C-terminal domains. The structures of three *Acinetobacter baumannii* PilA proteins, $PilA^{ACICU}$ (gold), $PilA^{AB5075}$ (magenta) and $PilA^{BIDMC57}$ (blue) are superimposed on structural similar pilin protein homologues from (respectively) *P. aeruginosa* K, *P. aeruginosa* 1244 and group A *D. nodosus*.

```
CLUSTAL O(1.2.4) multiple sequence alignment


ATCC_19606      FTLIELMIVVAIIGILAAIAIPAYQNYIAKSQASEAFTLADGLKTTINTNLQAGTCFAGG 60
M2              FTLIELMIVVAIIGILAAIAIPAYQNYIAKSQASEAFTLADGLKTTINTNLQAGTCFAGG 60
                ************************************************************

ATCC_19606      ATAATAADQVAGKYGDAEIGGTAPNCTITYTFKSSGVSTKLTSKQIVMNVSETGILTKNS 120
M2              ATAVTAADKVSGKYGDAEIGGTAPNCTITYTFKSSGVSNKLTSTKIVMNVSETGILTKNS 120
                ***.****:*:***************************.****.:***************

ATCC_19606      STNAPAELLPQSFTAS  136
M2              GTDTPVELLPQSFVAS  136
                .*::*.*******.**
```

**Supplementary Figure 2:** Clustal Omega alignment of mature PilA amino acid sequences from *Acinetobacter baumannii* ATCC 19606 (top) and *Acinetobacter nosocomialis* M2 (bottom)

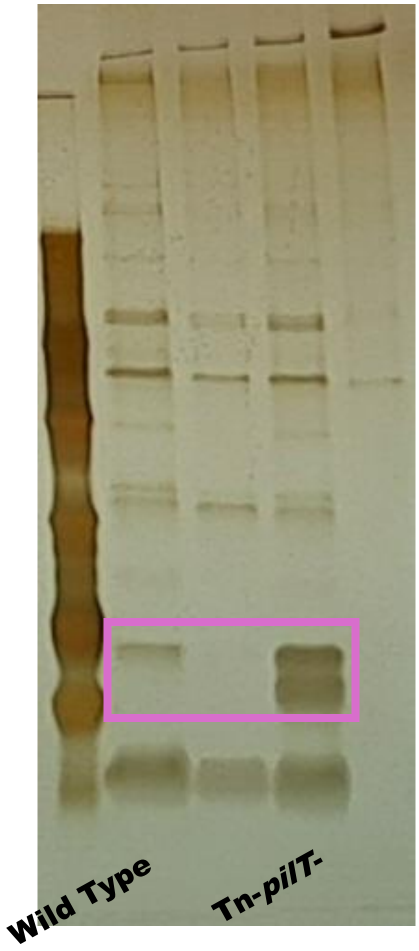


**Supplementary Figure 3:** Isolation of type IV pili from *A. baumannii* AB5075-UW.. Silver-stained gels are shown for pilus preparations (sheared and precipitated as described in Methods) for *A. baumannii* AB5075-UW wild type and Tn-*pilT*-.

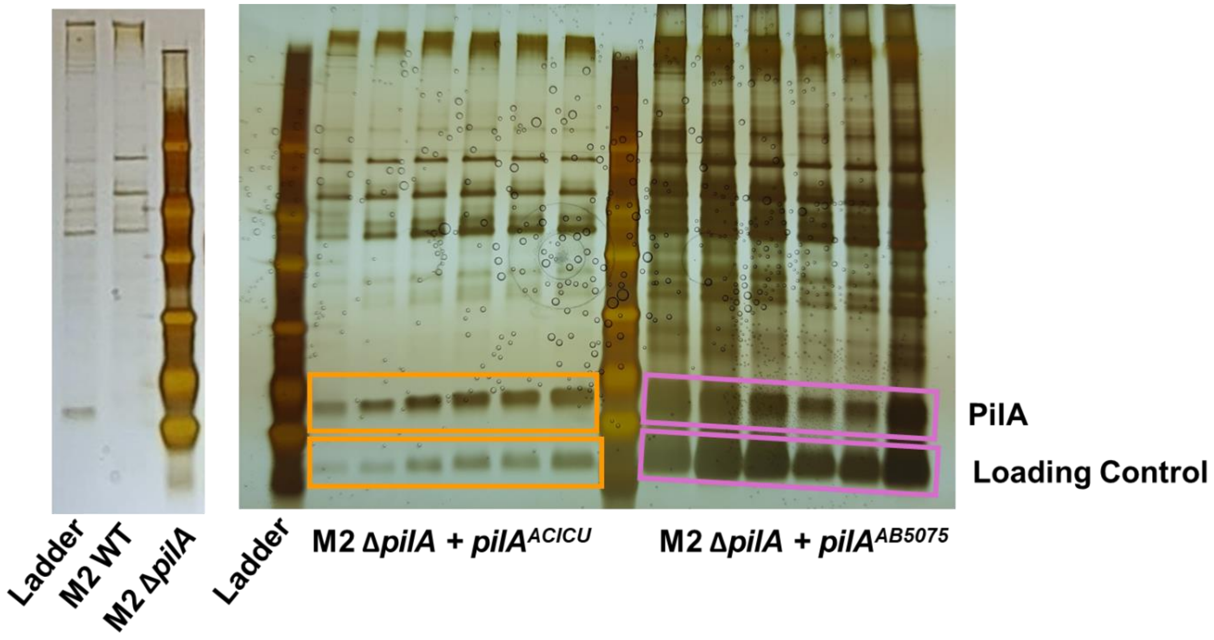


**Supplementary Figure 4:** Isolation of type IV pili from *A. nosocomialis* M2. Silver-stained gels are shown for pilus preparations (sheared and precipitated as described in Methods) for *A. nosocomialis* M2 wt, Δ*pilA* and complements with either *pilA*$^{ACICU}$ or *pilA*$^{AB5075}$.

| Construct | Vector | Sequence |
|---|---|---|
| *pilA*$^{ACICU}$ | pUCP20GM | ATGAATGCACAAAAAGGTTTTACATTAATCGAACT<br>CATGATCGTAGTTGCCATTATTGGTATTTTGGCTG<br>CGATTGCGATTCCTGCTTATCAAAACTACATTGCT<br>AAGTCACAAGTAAGTACTGGTTTAGCTGATATTAC<br>TGCTGGTAAGACAAACGCAGAAACTAAATTAGCAG<br>AAGGTTTAACTGCGGCATTAACTGATGTAGAAGCT<br>TTAGGCTTACAAAAATCTACGAATGCTTGTAGTAC<br>TATTACAACCAGTATCGGAACTAATGGTGCAAGTA<br>ATATTACTTGTACATTGAAAGGTACATCACAAATT<br>AATAGTAAAAAAATTGAATGGATCCGTGATGCAGA<br>TAATGCTACAAATGGTACGACAGGTGCTTGGCGCT<br>GTAAAACTGATGTAGCTGAAAACTTACGTCCTAAA<br>TCATGTGGTGCTTCTTAA |
| *pilA*$^{AB5075}$ | pUCP20GM | TTCACTCTGATTGAACTGATGATTGTTGTTGCAAT<br>TATTGGCATTTTAGCAGCTATTGCTATTCCACAAT<br>ATCAAACCTATATTGCAAAAAGCCAAGTTTCTCGT<br>GCTGTTAGTGAAAGCGGTTCTTTAAAAACAGTTAT<br>TGAAGATTGTCTGAATAATGGCAAAACCACAGTTG<br>GTGAAGCAGCTGGCGAATGCGCAATTGGTGCTACC<br>GGCTCAAATATTTTAGATGGTGCAGCTCAAAGTGG<br>CGAAACTTTAGCAGCTGGTACCGGCGTTCCACAAG<br>TTACATTAGCAAATACTGGTGCAGCTACCATTGTT<br>GCTACATTTGGCAATTCAGCAAGTACAGCTTTAAA<br>AAGCACTCCTACTACCGTTACCTGGACACGTACAA<br>CTGATGGTACTTGGACCTGTGAATCTACAGCAGCT<br>GAAAAATATAACTCTTCAGCTTGCCCTGCAGCT |
| *pilA*$^{ACICU-negative}$ | pUCP20GM | TTTACATTAATCGAACTCATGATCGTAGTTGCCAT<br>TATTGGTATTTTGGCTGCGATTGCGATTCCTGCTT<br>ATCAAAACTACATTGCTAAGTCACAAGTAAGTACT<br>GGTTTAGCTGATATTACTGCTGGTAAGACAAACGC<br>AGAAACTAAATTAGCAGAAGGTTTAACTGCGGCAT<br>TAACTGATGTAGAAGCTTTAGGCTTACAAAAATCT<br>ACGAATGCTTGTAGTACTATTACAACCAGTATCGG<br>AACTAATGGTGCAAGTAATATTACTTGTACATTGA<br>AAGGTACATCACAAATTAATAGTACAACTATTGAA<br>TGGATCCGTGATGCAGATAATGCTACAAATGGTAC<br>GACAGGTGCTTGGACATGTGAAACTGATGTAGCTG<br>AAAACTTAAACCCTAAATCATGTGGTGCTTCT |
| *pilA*$^{ACICU-loop\ swap}$ | pUCP20GM | ATGAATGCACAAAAAGGTTTTACATTAATCGAACT<br>CATGATCGTAGTTGCCATTATTGGTATTTTGGCTG<br>CGATTGCGATTCCTGCTTATCAAAACTACATTGCT<br>AAGTCACAAGTAAGTACTGGTTTAGCTGATATTAC<br>TGCTGGTAAGACAAACGCAGAAACTAAACTGAATA<br>ATGGCAAAACCACAGTTGGTGAAGCAGCTGGCGAA<br>TGCGCAATTGGTGCTACCGGCTCAAATGCTTGTAG<br>TACTATTACAACCAGTATCGGAACTAATGGTGCAA<br>GTAATATTACTTGTACATTGAAAGGTACATCACAA<br>ATTAATAGTAAAAAAATTGAATGGATCCGTGATGC<br>AGATAATGCTACAAATGGTACGACAGGTGCTTGGC<br>GCTGTAAAACTGATGTAGCTGAAAACTTACGTCCT<br>AAATCATGTGGTGCTTCTTAA |

**Supplementary Table 1:** Nucleotide sequences for *Acinetobacter baumannii pilA* genes *pilA*$^{ACICU}$, *pilA*$^{AB5075}$, *pilA*$^{ACICU-negative}$ and *pilA*$^{ACICU-loop\ swap}$.
